# Endothelial *Cept1* Promotes Post-Ischemic Angiogenesis in a *Pparα*-Dependent Fashion

**DOI:** 10.1101/2025.03.11.642511

**Authors:** Tariq J. Khan, Rodrigo Meade, Santiago Elizondo Benedetto, Larisa Belaygorod, Omar Saffaf, Brigida Rusconi, Fong-Fu Hsu, Sangeeta Adak, Batool Arif, Mohamed Zaghloul, Tiandao Li, Bo Zhang, Clay F. Semenkovich, Mohamed A. Zayed

## Abstract

**Background:** *Cept1* is essential for *de novo* phopholipogenesis and is impacted by diabetes. We previously demonstrated that conditional knockdown of *Cept1* in the endothelium leads to reduced hindlimb angiogenesis and tissue recovery. We hypothesized that *Cept1* may also be sufficient in promoting post-ischemic angiogenesis and recovery in the setting of diabetes.

**Methods:** CEPT1 content was evaluated in peripheral arteries of patients with peripheral arterial disease (PAD), and with or without diabetes. An endothelial cell (EC)-specific *Cept1* overexpression mouse model was developed (*Cept1^fl/fl^Cre*^+^) in adult C57BL6 mice. Murine aortae were harvested, for single-cell RNA sequencing (scRNA-seq), and unilateral hindlimb ischemia was used to evaluate angiogenesis in *Cept1^fl/fl^Cre*^+^ mice. Primary ECs were isolated and HUVECs transduced with *Cept1* cDNA were developed, and evaluated using molecular assays, *in vitro* functional assays, and mass spectrometry.

**Results:** In humans, arterial intima CEPT1 was elevated in the setting of PAD and diabetes, along with ACOX1, VEGF2R, p-Akt, and p-eNOS. In mice, scRNA-seq demonstrated that ECs with *Cept1* overexpression were enriched with wound healing, angiogenesis, sprouting, and cell migration pathways. Diabetic *Cept1^fl/fl^Cre^+^* mice had improved hind-limb perfusion and angiogenesis, and their aortic rings had increased *ex vivo* capillary sprouting. *Cept1* overexpression in ECs significantly increased migration, tubule formation, and proliferation as predicted by scRNA-seq. *Cept1* overexpression in ECs led to increased *Pparα, Acox1, Vegfa,* and *Vegf2r*. Similarly, treatment with si*Pparα*, and inhibitors for PPARα (GW6471), VEGFR2 (ZM323881), Akt (LY294002), and eNOS (L-*NAME*) abrogated CEPT1-induced EC migration.

**Conclusions:** *Cept1* overexpression promotes EC function and post-ischemic recovery. The impact of CEPT1 on ECs is at least in part dependent on p-Akt/p-eNOS angiogenic signaling and PPARα. Since CEPT1 is elevated in diseased human peripheral arterial tissue, these findings suggest that CEPT1 may be playing an important compensatory role in vascular recovery and reperfusion following ischemic injury in the setting diabetes.

**Highlights:**

- CEPT1 content is higher in the peripheral arteries of individuals with peripheral arterial disease (PAD) and type 2 diabetes.
- *Cept1* over expression induces endothelial cell activation and function and enhances post-ischemia angiogenesis *in vivo*.
- CEPT1 induces endothelial pAkt/p-eNOS signaling and VEGF-A production in a PPARα dependent fashion.
- CEPT1 may be an important regenerative signal that is increased in the peripheral arteries in the setting of PAD.

**Graphical Abstract:** 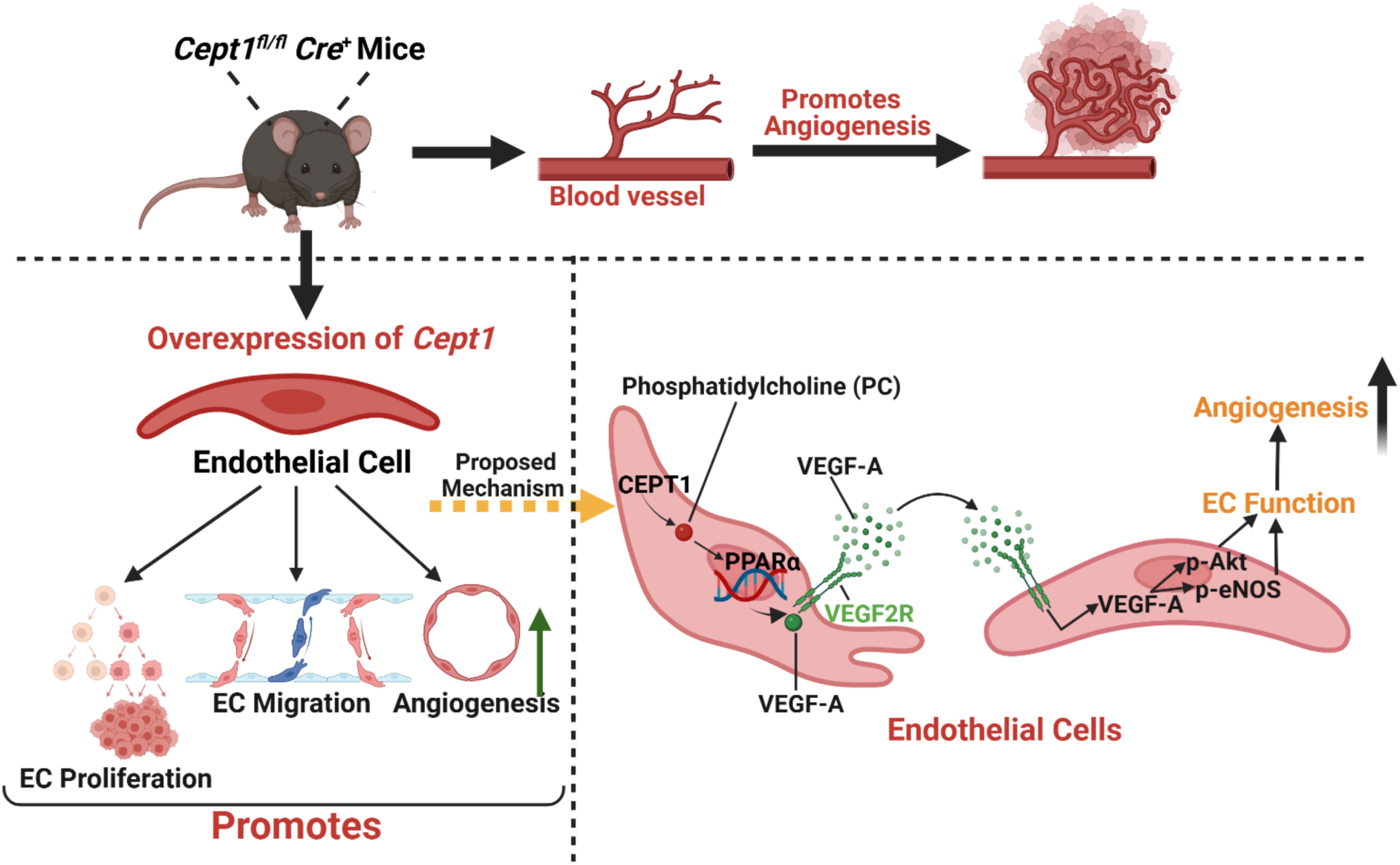

## Introduction

Peripheral arterial disease (PAD) is a common vascular pathology that impacts lower extremity function and is associated with high morbidity.^1^ This results from luminal narrowing in the peripheral arteries leading to reduced oxygenated blood flow to the affected limb, causing increased risk of wounds, tissue loss, and amputation.^1^ Globally, more than 200 million individuals suffer from PAD, particularly in individuals with diabetes who have higher prevalence and risk of complications as a result of PAD.^1^ In this setting, compensatory molecular mechanisms that can help restore arterial blood flow via angiogenic signaling can be vitally important in alleviating chronic manifestations of PAD in high-risk diabetes patients.^2^ To date, there are only a few FDA pharmacological interventions that are aimed to manage the progression of chronic PAD in the setting of diabetes.^3,4^ There continues to be a persistent need for the identification of important molecular targets that can guide future drug discovery in this disease.

*De novo* phospholipogenesis is a highly conserved process in mammals that endogenously produces phospholipids using excess dietary carbohydrates and proteins.^5,6^ The Kennedy Pathway, which is primarily responsible for *de novo* phospholipogenesis, requires choline-ethanolamine phosphotransferase 1 (CEPT1), a 47-kDa membrane-bound protein that is expressed by the *Cept1* gene.^7,8^ CEPT1 is responsible for the synthesis of >50% of phospholipids, specifically phosphatidylcholine (PC) and phosphatidylethanolamine (PE).^8,9^ PC and PE play critical functions in maintaining plasma membranes, lipoprotein formation, and intracellular signaling responsible for DNA transcription and protein translation.^10–12^ Recent evidence shows that metabolic disorders such as diabetes can lead to altered tissue phospholipogenesis ^13–16^, and that CEPT1 is elevated in the arterial intima of individuals with diabetes who are suffering from peripheral arterial disease (PAD) ^17^. Conditional knockdown of *Cept1* in the endothelium reduces ischemia-induced tissue recovery in a Peroxisome Proliferator-Activated Receptor α (PPARα)-dependent manner.^17^

PPARα is a transcription factor that is activated by endogenously produced ligands, such as the CEPT1-derived phospholipid derivative PC16:0/18:1,^18^ and impacts intracellular lipid metabolism through enzymes such as acyl-CoA oxidase (ACOX1) and carnitine palmitoyl transferase (CPT1).^19,20^ While it is known that PPARα can induce neovascularization through vascular endothelial growth factor A (VEGF-A) signaling,^17,21,22^ it is unknown whether PPARα activation via CEPT1 can impact angiogenesis and endothelial cell (EC) function. To evaluate this, we developed a conditional EC-specific *Cept1* overexpression (*Cept1^fl/fl^ Cre^+^*) mouse model and evaluated whether *Cept1* can sufficiently impact EC function and angiogenesis *in vivo* and *in vitro*.

## Methods

### Human Arterial Tissue Analysis

Human peripheral arterial tissue was obtained from the Washington University Vascular Biobank, which is institutional review board (IRB)-approved. All patients who participate in the biobank provided written informed consent prior to participation. Peripheral arterial specimens were obtained from 12 patients with and without chronic type 2 diabetes (T2D) who were undergoing lower extremity amputation due to advanced non-salvageable peripheral arterial disease (PAD). Tibial arterial specimens were harvested *en bloc* from the lower legs immediately after amputation in the operating room, placed in a cold saline solution, maintained on ice, and immediately transferred to the laboratory for further analysis. Arterial tissue was subdivided into maximally (Max) and minimally (Min) diseased arterial segments as previously described.^23^

### Animal Regulations

All housing, breeding, and experimental procedures involving mice were conducted by national guidelines and regulations and approved by the Washington University institutional animal care and use committee (IACUC).

### Generation of Mouse Models

A conditional EC-specific *Cept1* overexpression mouse model was engineered by breeding *Cept1^fl/fl^* mice (Figure S1) with *VE-cadherin*-*Cre*-*ERT2* mice.^24^ Male littermates at least 8 weeks of age received tamoxifen (TMX-diluted with sunflower oil) treatment (five daily i.p. injections at 1 mg per mouse) to induce *Cre* expression and conditional EC-specific *Cept1* gene over-expression. Mice were allowed to recover for 5 days post TMX injections before initiating *in vivo* experimental protocols.

### Digestion of the aorta and single-cell sorting

Male mice were euthanized, and the thoracic and abdominal aorta were surgically micro-dissected from the aortic arch to the distal aortic bifurcation in the pelvis. Aortic tissue (n = 3 per condition) was minced into small pieces using a scalpel and transferred into a glass vial with 1.5 mL of cold digestion buffer (125U/ml Collagenase XI, 60U/ml Hyaluronidase type 1-s, 60U/ml DNase I and 450U/ml Collagenase type I. Tissue was further digested in a shaking (100 rpm) 37 °C water bath for 1 hour. The resulting cell suspension was strained through 70μM cell strainer and treated with ACK lysis buffer to lyse red blood cells. Single-cell suspensions then used for subsequent single-cell sorting.

### Single Cell RNA-Sequencing and Bioinformatics Analysis

Transcriptional profiling of *Cept1^fl/fl^ Cre*^−^ and *Cept1^fl/fl^ Cre*^+^ mouse whole aortae was performed using droplet-based massively parallel 10X genomic single-cell RNA sequencing (scRNA-seq) 3’ v3. Single cells for each genotype were individually barcoded and subjected to sequencing, generating Fastq files. Fastq files were then processed with Cellranger to yield a unique molecular identifier (UMI) matrix, filtering out low-quality barcodes and mapping reads to the mouse genome (mm10), resulting in approximately 15,000 cells and 2,000 genes per cell per group. Datasets from both genotypes were combined using the Cellranger aggr pipeline to equalize the average read depth between samples. Gene expression from both samples were filtered, normalized, and clustered using the R package Seurat. To filter out multiplets, low-quality cells, and empty droplets, features were normalized to include log2 counts between 9.3-14.5, UMI log2 count within 8.5-11, and mitochondrial genes below 15%. Principal component analysis (PCA) was performed to reduce dataset dimensionality, and t-distributed stochastic neighbor embedding (t-SNE) was used to visualize principal components, focusing on the first two dimensions. Unbiased clustering and marker identification defined each cell types, and differentially expressed genes between clusters were identified for pathway analysis in Metascape. Bonferroni-adjusted p-values were used to determine significance at an FDR adjusted p value of <0.05. Genes involved in pathways analysis were represented using module scores based on average expression calculation within a specific cluster.^25^

### Isolation of Primary EC

Liver and lung organs were harvested from 6–8-week-old *Cept1^fl/fl^ Cre*^−^ and *Cept1^fl/fl^ Cre*^+^ mice that underwent TMX-induced *VE*-*cadherin*-*Cre*-*ERT2* expression. Organs were removed from the mediastinum and abdomen *en bloc*, and prepared for mouse EC isolation as previously described.^26^ Briefly, organs were gently minced, digested with Collagenase I (Worthington-biochemical), and strained. The resulting cell suspension underwent positive cell sorting using PECAM-1 (BD Biosciences, San Jose, CA) anti-rat IgG-conjugated magnetic beads (Invitrogen). Isolated cells were then plated in tissue culture flasks and cultured using endothelial growth media-2 (EGM-2; Lonza Bioscience). Liver and lung-derived ECs were then purified with a second round of positive cell sorting using PECAM-1 coated magnetic beads. The remaining ECs were then cultured for *in vitro* experiments and were not used beyond 3 primary passages. Mouse ECs were isolated using isolation media [i.e., high glucose (4.5 g/L glucose) DMEM with 20% FBS, 20 units/mL penicillin/streptomycin antibiotic (Gibco), and 20 units/L heparin (Sigma)]. Cells were maintained in culture using EBM culture media according to manufacturer instructions (Cambrex).

### Monolayer Wound Healing Assay

Migration of mouse liver and lung-derived ECs as well as Human Umbilical Vein EC (obtained from ATCC) was evaluated using a monolayer wound healing assay as previously described.^26,27^ inhibitors including GW6471 (PPARα inhibitor-10μM), ZM323881 (VEGF2R inhibitor-80nM), LY294002 (Akt inhibitor-30μM), and L-*NAME* (eNOS inhibitor-0.5μM) were used toassess responses. EC monolayer scratches were made using a 200μL micropipette tip and were serially evaluated at 0, 6, and 16 hours. The percentage of wound area closure was measured using ImageJ software. Each condition was repeated in triplicate.^17^

### HUVEC Culture, Transduction, and Transfection

HUVECs were cultured using EGM-2 media according to the manufacturer’s instructions (Cambrex) and studied within 3 primary passages. HUVECs were transduced to overexpress human *Cept1* with lentiviral ORF technology and control lentiviral ORF particles were used to generate a control cell line for the overexpression system (Pan02^OE-CTL^; Origene NM_001007794 and PS100093V; Origene Technology). All transductions were performed according to the manufacturer’s instructions, and as previously described.^28^ Multiplicity of Infection (MOI) is the number of transducing lentiviral particles per cell and we used MOI5 for the viral transduction.^28^ The transduction efficiency was confirmed by quantitative real-time PCR analysis and Western blotting. HUVECs were transfected with siRNA-*Pparα* (10nmol/L; AM16708-Thermo Fischer Scientific) and non-targeting siRNA (Negative Control #1 siRNA, Thermo Fisher Scientific) using Lipofectamine RNAiMAX Reagent (Thermo Fisher Scientific) according to the manufacturer’s recommendations.

### Tube Formation Assay

Mouse primary ECs and HUVECs were evaluated for tubule formation using a 96-well culture format as previously described.^26^ Approximately 1×10^4^ mouse primary ECs and *Cept1*-transduced HUVECs were seeded per well on the growth factor-reduced Matrigel (biotechne; RD systems) and then incubated in the presence of growth media (EGM2) or basal media. Three random 10× magnification images were collected for each condition at baseline and 6 hours post-incubation. Each assay was performed in triplicate and quantified using ImageJ.^17^

### Cell Proliferation and Viability Assay

Primary EC and HUVEC proliferation were evaluated using a BrdU incorporation assay, and viability was evaluated using an anti-DNA/histone ELISA. In a 96-well culture format, 1×10^4^ liver and lung primary ECs and *Cept1*-transduced HUVECs were seeded per well, followed by serum starvation for at least 6 hours. Growth media was then added, and proliferation and death were independently evaluated after 24 hours according to the manufacturer instructions (Roche, Indianapolis, IN) using a multi-well spectrophotometer at 450 nm as previously described.^26,27^ Each assay was repeated in triplicate.

### Mouse Aortic Ring Assay

Aortic ring assays were performed using aortic specimens harvested from *Cept1^fl/fl^ Cre*^−^ and *Cept1^fl/fl^ Cre*^+^ mice.^29^ Briefly, the descending thoracic aorta was surgically harvested *en bloc*, and 1-mm long aortic rings were embedded in growth factor-reduced Matrigel (354234, Corning). The aortic rings were then cultured in Gibco™ Opti-MEM™ I (Thermo Fisher Scientific) supplemented with 10% FBS in a humidified 37°C, 5% CO2 incubator for 6 days. NIH ImageJ was used to analyze the length of aortic sprouts (n=6).^17^

### Murine Hind-Limb Ischemia

Aged matched (7-week-old) *Cept1^fl/fl^ Cre*^−^ and *Cept1^fl/fl^ Cre*^+^ mice were maintained on a regular diet and pre-treated with or without streptozotocin (STZ; 0.1 mg/g of body weight administered i.p. once daily over five consecutive days) to induce hyperglycemia and a diabetes-like phenotype.^17^ At 10 days after *Cept1* overexpression, unilateral hind-limb ischemia (HLI) was performed as described.^17^ At 0, 3, 7, 14 and 21 days post-HLI, mouse hind-limb Doppler perfusion was performed. At 21 days post-ischemia, the gastrocnemius muscle was harvested *en bloc* and evaluated using muscle fiber diameter/size and microvessel density as described.^17^ As previously described, ischemic limb and appearance scores were also determined with some modifications.^26^ Limb appearance was independently graded on a scale of 0–4 (4 = digit/foot autoamputation, 3 = severe discoloration/gangrene, 2 = moderate discoloration, 1 = mild discoloration, 0 = normal appearance). Limb use was independently graded on a scale of 0–3 (3 = foot-dragging, 2 = no foot-dragging but no plantar flexion, 1 = abnormal plantar flexion, 0 = normal foot and leg function). Twenty-one days after femoral artery ligation, mice were administered a cocktail of ketamine (80 mg/kg) and xylazine (10 mg/kg). Cold PBS with heparin (10 units/mL) was injected into the gastrocnemius muscles before harvest for subsequent protein analysis or fixed and paraffin embedding.^17^ For each muscle specimen, representative 10-μm sections were obtained and stained with hematoxylin-eosin or labeled with Alexa Fluor 594–conjugated *Griffonia simplicifolia* isolectin-1-B4 1:100 (Invitrogen). Five 20× images were collected for each section from each mouse genotype. Using ImageJ, muscle atrophy and microvascular capillary density were quantified as previously described.^17^

### Tissue Lysis, Western Blotting and Immunostaining

Cells and tissues were lysed for protein analysis. ∼50 mg of arterial tissue was homogenized in cold RIPA Lysis Buffer System (sc-24948; Santa Cruz Biotechnology) containing Halt Protease and Phosphatase Inhibitor Cocktail (78442; Thermo Fisher Scientific). Total protein from mouse ECs, HUVECs, and carotid arteries was determined using a Bradford protein assay, loaded onto SDS-PAGE gel, and transferred to polyvinylidene fluoride membranes for Western blotting. Proteins for western blot and immunostaining were detected with rabbit anti-CEPT1 (bs-12284R; Bioss USA), mouse anti-PPARα (66826-1-1g; Proteintech), rabbit anti-ACOX1 (10957-1-AP; Proteintech), rabbit anti-VEGF-A (bs-1665R; Bioss USA), rabbit anti-VEGF2r (#2472; Cell Signaling), rabbit anti-eNOS (PA1-037; Thermo Fisher), rabbit anti-p-eNOS (MA5-14957; Thermo Fisher), rabbit anti-Akt (MA5-14916; Thermo Fisher), rabbit anti-p-Akt (#9271; Cell Signaling), and mouse anti-CD31 (sc-376764; Santa Cruz Biotechnology, Inc). Rabbit anti-GAPDH (ab2910; Abcam Biotechnology) was used for Western blot loading controls. Band densitometry analysis was performed using ImageJ software.^26,27^ Band densities were averaged across triplicate blots and expressed as ratios relative to protein loading control or non-phosphorylated total protein. Immunostaining was performed using primary antibodies and primary antibodies was detected with a secondary antibody donkey anti-rabbit IgG labeled with Alexa Fluor 555 1:400 (A31572; Thermo Fisher Scientific), followed by DAPI staining. Imaging assessments were performed using a Leica THUNDER Imaging Systems microscope, and staining was quantified using ImageJ software.^30^

### Real-time PCR

RNA was purified as previously described.^23^ Target mRNA was quantified by real-time PCR using specified primer sets (designed to span introns, not react with genomic DNA, and validated with quantitative PCR melting curve analysis) and Applied Biosystems PowerUp SYBR Green Master Mix (A25742; Thermo Fischer Scientific). Samples were evaluated using the 7500 Fast Real-Time PCR System (Applied Biosystems) and analyzed using Fast System Software (Applied Biosystems). Threshold cycle (Ct) values were normalized to *Gapdh* mRNA and expressed as fold change relative to the mean Ct value of the control group using the Δ Ct method.

### Electrospray Ionization Mass Spectrometry

Lipid extracts from ECs were analyzed by direct injection electrospray ionization mass spectrometry using a Thermo Vantage triple-quadruple mass spectrometer (San Jose, CA) and an Accela 1250 UPLC system operated via the Xcalibur operating system. Structures for all phospholipids were identified using a multiple-stage linear ion trap that was operated at a low-energy collision-induced dissociation and high-resolution mass spectrometry. Phospholipid structural assignments, phospholipid content, and mass spectrometry–derived lipid mass spectrum for EC samples were made as previously described.^17^

### Statistical Analysis

Tukey multiple comparison two-way ANOVA was used to evaluate differences in EC tubule sprout formation and migration. Sidak multiple comparison one-way ANOVA was used to evaluate differences in hind-paw Doppler perfusion. Unpaired two-tailed Student *t* test was used to evaluate differences in gene expression in ECs and protein content. Unpaired two-tailed Student *t* test was also used to evaluate differences in PCs and PEs in mouse ECs. Unpaired two-tailed Student *t* test with Welch correction was used to evaluate differences in hind-limb Doppler perfusion, relative muscle fiber size, and relative microvessel density. We considered p < 0.05 to be significant. Error was presented as SEM.

### Data and Resource Availability

The data sets generated and/or analyzed during the current study are available from the corresponding author upon reasonable request.

## Results

### CEPT1 content in variably diseased peripheral arterial segments

Peripheral arterial segments were analyzed from 24 human subjects. This included 3 healthy organ donor patients and 21 patients with advanced non-salvageable PAD requiring major lower extremity amputation. A large proportion of patients with PAD had cardiovascular risk factors, including hyperlipidemia (74.2%), hypertension (90.3%), former/current smoking (83.9%), and type 2 diabetes (34.7%). Compared to healthy controls, CEPT1 protein content was significantly elevated in maximally (Max)-diseased versus minimally (Min)-diseased arterial segments, and in patients with T2D (Figure 1B-E). Immunostaining also demonstrated significantly higher CEPT1, ACOX1, VEGF2R, p-Akt, and p-eNOS protein content in ECs of the Max-diseased arterial segments of patients with T2D compared to healthy controls (Figure 1F-J).

**Figure 1:**
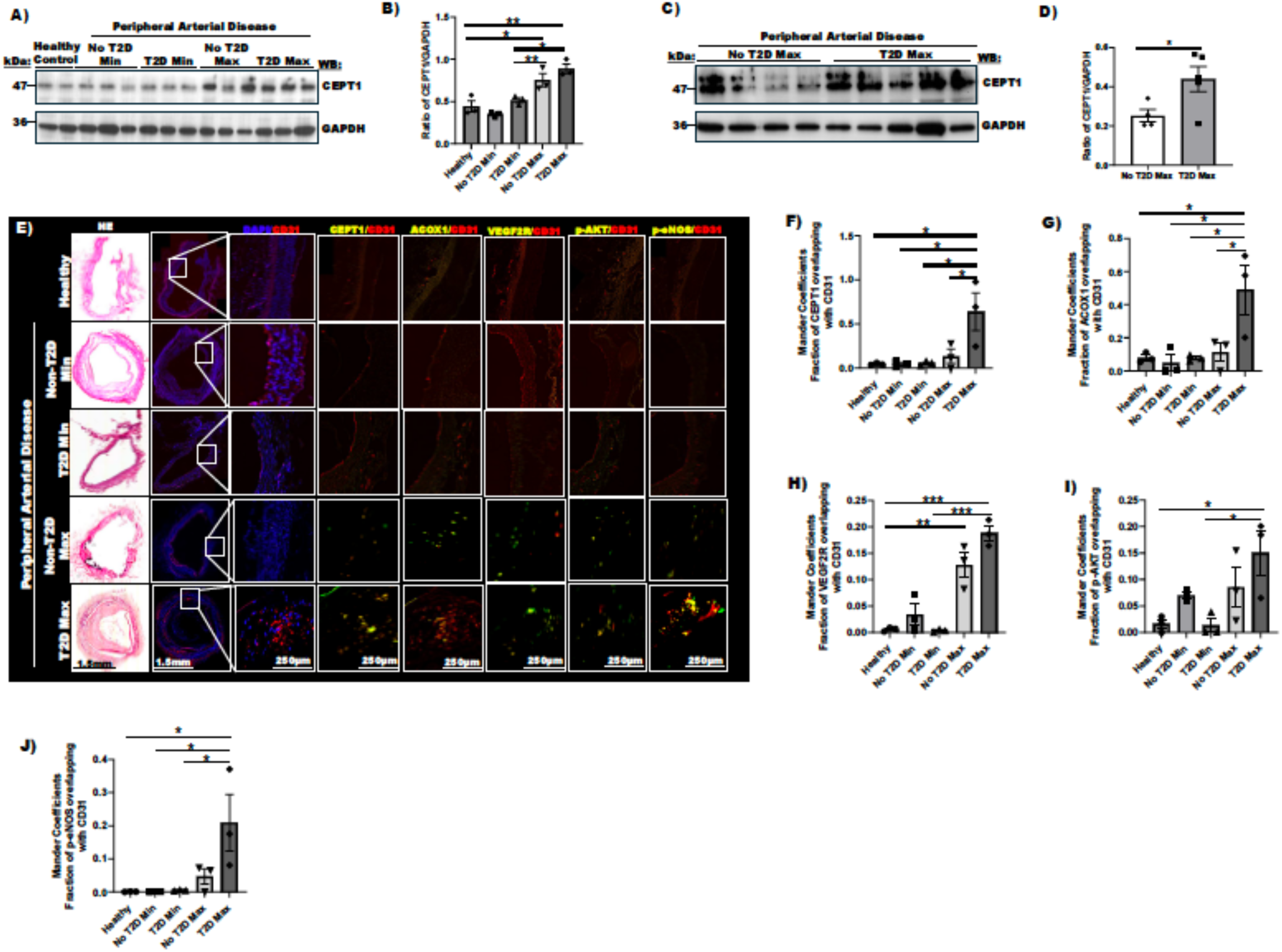
CEPT1 is elevated in the arterial segment of diseased peripheral arteries. A) Min and Max diseased peripheral arterial segments were procured from the lower extremities of patients with severe PAD, and with or without T2D. Peripheral arterial segments were also harvested from health organ donors as controls. B-E) Western blot of CEPT1 in healthy, non-T2D Min, T2D Min, non-T2D Max, and T2D Max-diseased arterial segments was quantified relative to GAPDH (n=3-8). F-J) Immunostaining of arterial segments for CEPT1, ACOX1, VEGF2R, p-Akt, and p-eNOS that colocalized with CD31. Colocalization between two channels (fractions of CEPT1, ACOX1, VEGF2R, p-Akt, and p-eNOS colocalized with CD31) were expressed by Manders coefficients. Data are presented as mean ± SEM. * p<0.05; ** p<0.01; *** p<0.001. *P* values were from 1-way ANOVA and unpaired Student *t*-test.

### Murine EC-specific *Cept1* overexpression leads to increased angiogenic gene signaling

Based on observations in human peripheral arterial samples we developed a mouse model of EC-specific inducible *Cept1* overexpression (*Cept1^fl/fl^ Cre*^+^), and performed gene expression and pathway analysis using single-cell RNA sequencing (scRNA-seq; Fig. 2A and Fig. S1). Aortas isolated from *Cept1^fl/fl^ Cre*^-^ (control) and *Cept1^fl/fl^ Cre*^+^ mice were processed for droplet scRNA-seq (Fig. 2A). After quality control of droplet scRNA-seq data, removal of immune cells was performed. Remaining clusters were classified based on the expression of canonical markers for ECs, fibroblasts, and vascular smooth muscle cells (VSMC; Fig. 2B&C and Fig S2A). To investigate the angiogenic role of conditional overexpression of *Cept1* in ECs, we further sub-clustered the ECs based on previously described canonical markers. EC1: *Vcam1, Clu, Gkn3, Eln*; EC2: *Cd36, Fabp4, Lpl, Gpihbp1, Flt1*; and EC3: *Lyve1* (Fig. S2A-C).^31^ *Cept1^fl/fl^ Cre*^+^ mice demonstrated an increased relative percentage of all EC clusters, mainly driven by the EC1 sub-cluster, compared to controls (Fig. 2D). *Cept1^fl/fl^ Cre*^+^ mice showed significantly elevated *Cept1* gene expression within the EC cluster compared to controls (p<0.001; Fig. S2D), again driven by the EC1 sub-cluster (p<0.001; Fig. 2D). *Cept1* overexpression in the EC1 sub-cluster resulted in a significant upregulation of genes involved in pathways associated with multiple angiogenic mechanisms, such as wound healing, sprouting angiogenesis, and cell migration (Fig. 2E-H and Fig. S2E).

**Figure 2:**
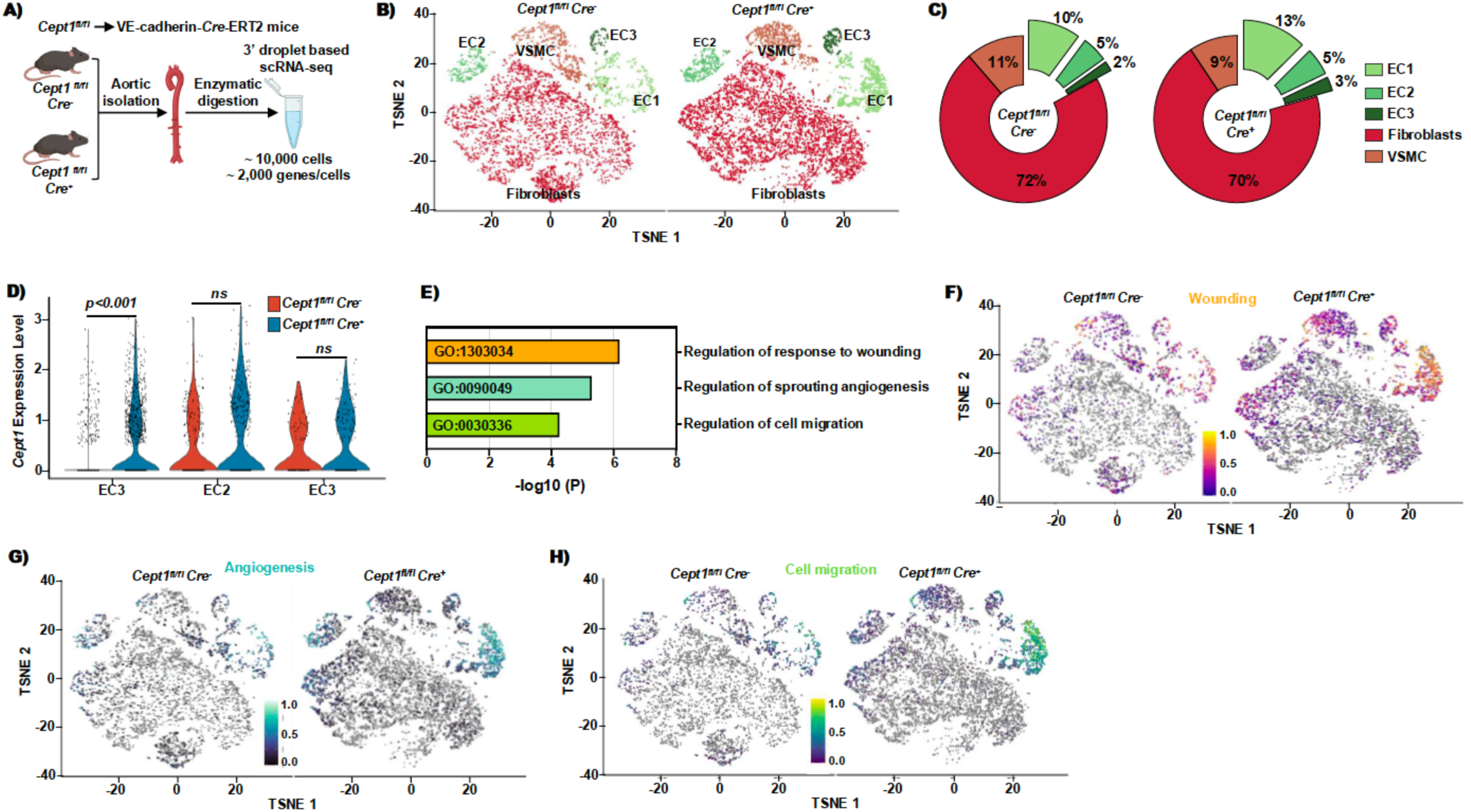
ScRNA-seq analysis of aortic tissue from mice with conditional EC-specific overexpression of *Cept1*. A) Experimental design of aortic cell isolation for scRNA-seq analysis. B) t-SNE of EC and stromal cells separated by genotypes. C) Pie-chart representing the relative abundance of each cluster. D) Violin plot of *Cept1* gene expression in EC sub-cluster. E) Pathway analysis of Differential Expressed Gene (DEG) between *Cept1* overexpression and control in EC1 sub-cluster (FDR adjusted p value of <0.05, log (FC) > 0.1 and pct.1 >0.6). F-H) Module score from significantly upregulated pathways.

### EC-specific *Cept1* overexpression enhances post-ischemic angiogenesis and perfusion

To validate the upregulation of angogensis pathways identfied in the sc-RNA Seq dataset we used an animal model of peripheral arterial disease. We induced hindlimb ischemia via unilateral femoral artery ligation ^32,33^ in *Cept1^fl/fl^ Cre^−^ and Cept1^fl/fl^ Cre^+^* mice. During the experiment, mice were also treated with STZ to induce a diabetes-like phenotype and were maintained on a regular diet (Figure 3A and Figure S3A). Hindlimb Doppler perfusion in *Cept1^fl/fl^ Cre^−^* and *Cept1^fl/fl^ Cre^+^* mice not treated with STZ demonstrated mild improvement in high paw perfusion at day 14 following ischemia (Figure S3B-D). Among mice treated with STZ, *Cept1^fl/fl^ Cre*^+^ mice demonstrated a significant improvement in gastrocnemius perfusion at days 3, 7, 14, and 21 (p<0.05; Figure 3B-F). Moreover, gastrocnemius muscle microvascular density was elevated in STZ-treated *Cept1^fl/fl^ Cre*^+^ mice (p<0.05; Figure 3G and H), and muscle fiber size was also increased in the *Cept1^fl/fl^ Cre*^+^ mice (p<0.05; Figure 3G and I) Non-significant improvement was observed in the non-STZ group (Figure S4A-E).

**Figure 3:**
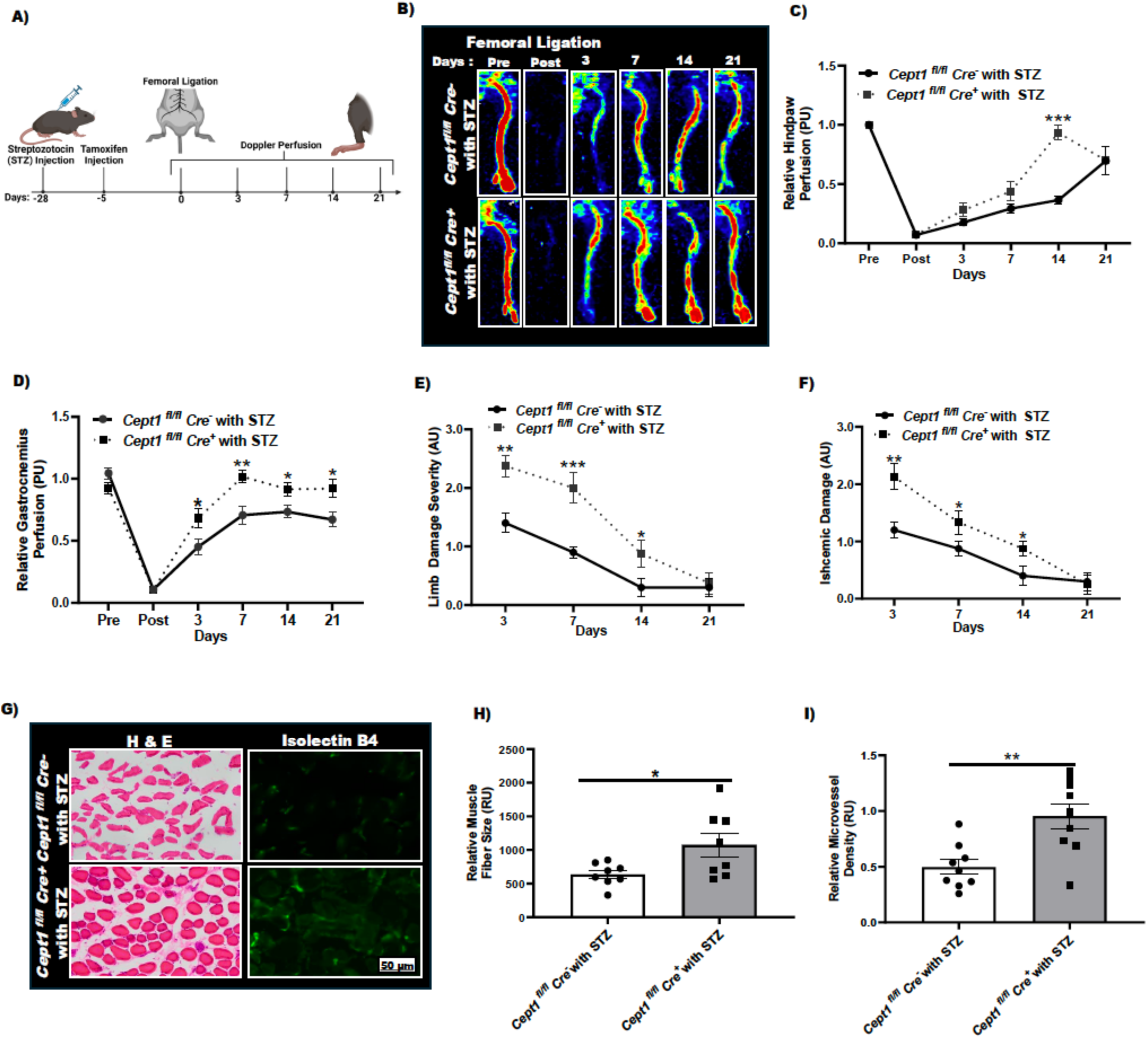
*Cept1* overexpression improves blood flow recovery after HLI. A) Graphical representation of the experiment design. B) Representative Doppler perfusion of HLI of *Cept1^fl/fl^ Cre*^−^ and *Cept1^fl/fl^ Cre*^+^ mice (n=8 mice) treated with STZ. C-D) Quantitative graphical representation of Doppler perfusion of the hind-paw and gastrocnemius muscle. E-F) Limb and ischemic damage severity were evaluated on days 3, 7, 14, and 21. G) Representative images showing HE and Isolectin B4 (IB4) staining in the gastrocnemius muscle at 21 days after artery ligation (n=8 mice). Scale bar =50 μm. H-I) Quantification of relative microvessel density and relative muscle fiber area (n=8 mice). Data are presented as mean ± SEM. * p<0.05; ** p<0.01; *** p<0.001. p values were from unpaired Student *t*-test.

### *Cept1* promotes EC migration, proliferation, and function

Given the impact of *Cept1* overexpression on hind-limb recovering, we next evaluated whether *Cept1* is sufficient in promoting EC function. Primary ECs isolated from *Cept1^fl/fl^ Cre*^+^ demonstrated significantly higher cellular proliferation (p<0.05; Figure 4A and B) and increased migration (p<0.05; Figure 4C-E). Similarly, aortic rings isolated from *Cept1^fl/fl^ Cre*^+^ mice demonstrated increased ring sprouting (p<0.05; Figure 4F-H). Primary ECs from *Cept1^fl/fl^ Cre*^+^ mice demonstrated a higher density of tubule formations on matrigel (p<0.05; Figure 4I and J). HUVEC transfected with *Cept1* cDNA to express higher *Cept1* (p<0.05; Figure 4K), also demonstrated higher cellular proliferation, cell migration, and tubule formation (p<0.05; Figure 4L-R). These findings suggest that *Cept1* overexpression promotes EC function.

**Figure 4:**
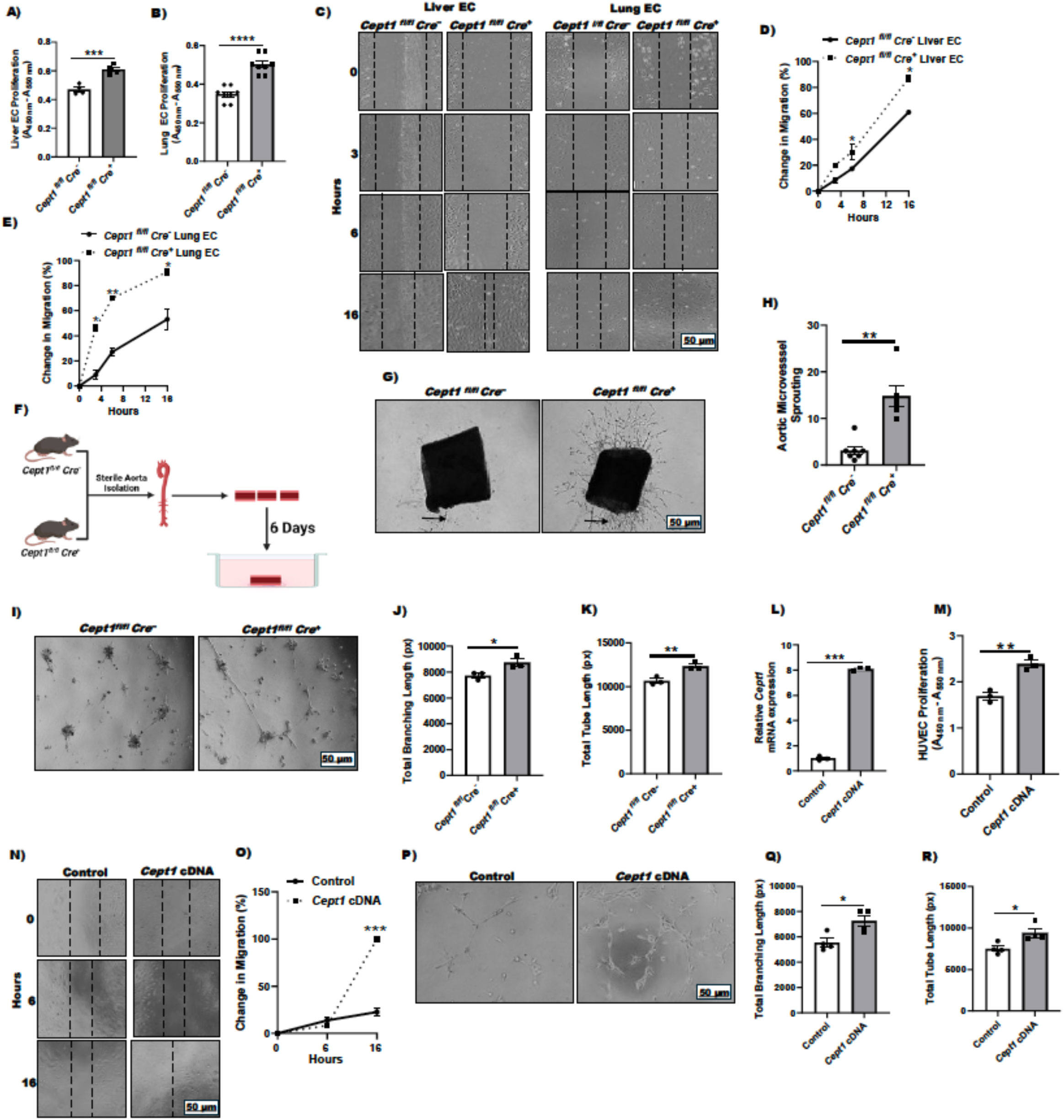
CEPT1 promotes EC function. A-B) Cell proliferation of the EC isolated from the liver and lung of *Cept1^fl/fl^ Cre*^−^ and *Cept1^fl/fl^ Cre*^+^ (n=3). C) Migration of cells was assessed using scratch assay from EC from the liver and lung of *Cept1^fl/fl^ Cre*^−^ and *Cept1^fl/fl^ Cre*^+^ (n=3 per condition). D-F) Quantitative representations of percent change in migration at different time points (t= 0, 3, 6, and 16h; n=3). F-H) Aortic ring assay to evaluate the growth of sprouts from the aortic rings of *Cept1^fl/fl^ Cre*^−^ and *Cept1^fl/fl^ Cre*^+^ were represented and quantitively analyzed at day 6 (n=3). I-K) Quantitative analysis of percent of tube formation of cells (n=3). L) Efficiency of *Cept1* lentiviral transduction was confirmed using RT-PCR (n=3). M) Proliferation of HUVEC treated with *Cept1* lentiviral particles (n=3). N) Representative image of the change of cell migration from HUVEC treated with *Cept1* lentiviral particle and its control (t=0, 6, and 16h). O) Quantitative analysis of percent change of migration of cells (n=3). P-R) Representative image of the tube formation from HUVEC treated with *Cept1* lentiviral particle and its control. Quantitative analysis of percent of tube formation of cells (n=3). Data are presented as mean ± SEM. * p<0.05; ** p<0.01; *** p<0.001. p values were from unpaired Student *t*-test.

### *Cept1* promotes EC activation and angiogenic signaling

Relative to primary ECs isolated from *Cept1^fl/fl^ Cre*^−^ mice, ECs isolated from *Cept1^fl/fl^ Cre*^+^ mice demonstrated significantly increased CEPT1, PPARα, ACOX1, VEGF2R, p-AKT, and p-eNOS (p<0.05; Figure 5A and B). Similarly, we observed increased expression of *Cept1*, *Pparα, Acox1, Vegfa* and *Vegf2r* (p<0.05; Figure 5C). Additionally, we observed significantly higher content of aPC and pPE levels in EC-isolated from *Cept1^fl/fl^ Cre*^+^ mice (p<0.05; Figure S5). HUVECs transduced with *Cept1* lentivirus demonstrated increased CEPT1, PPARα, ACOX1, VEGF2R, p-AKT, and p-eNOS content relative to control (p<0.05; Figure 5D and E). Similarly, transduced HUVECs had significantly higher expression of *Pparα*, *Acox1, Vegfa,* and *Vegf2r* (p<0.05; Figure 5F).

**Figure 5:**
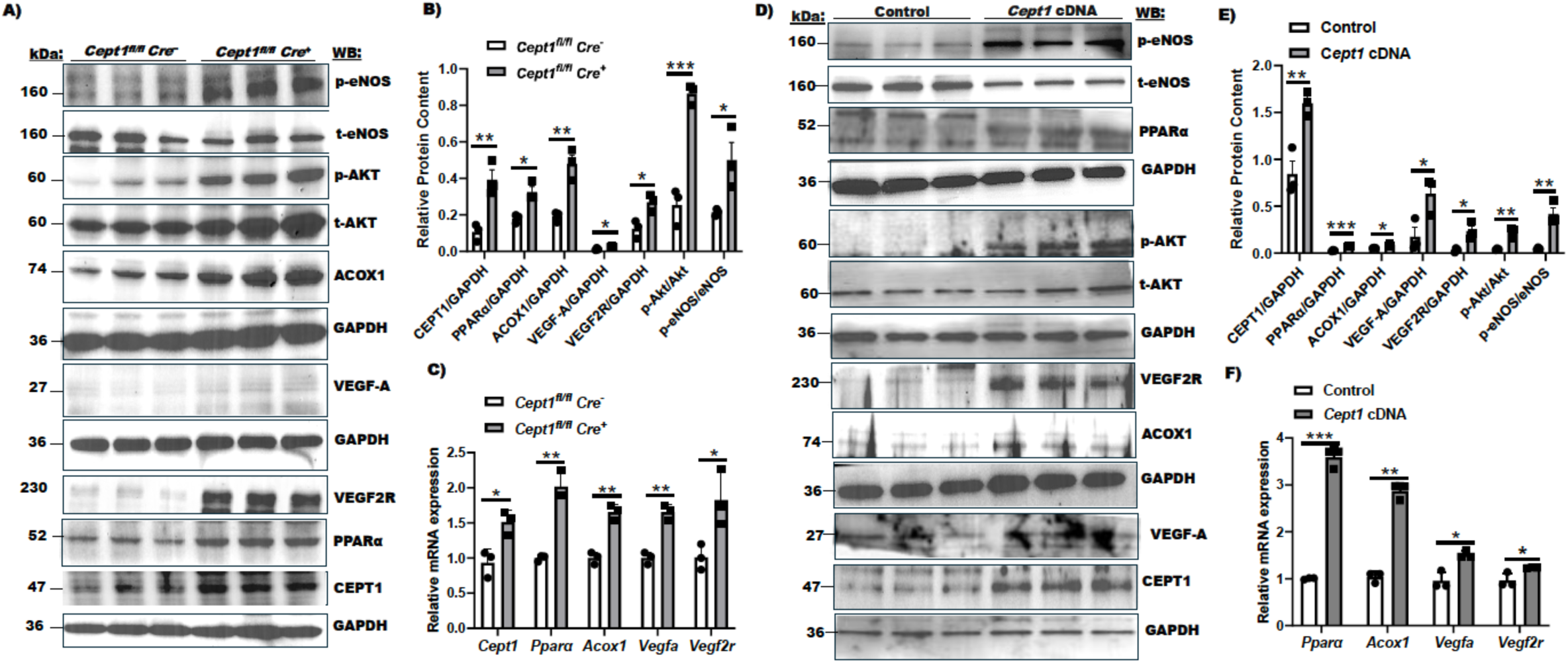
*Cept1* promotes the expression of angiogenic factors in ECs. A) Western blotting demonstrates that CEPT1 content is increased in ECs isolated from *Cept1^fl/fl^ Cre*^+^ mice. Representative blots of VEGF2R, PPAR*α*, ACOX1, p-Akt, t-Akt and p-eNOS, t-eNOS are also provided. B) Quantitative representation of protein content relative to GAPDH as a loading control (n=3). C) RT-PCR analysis demonstrates *Cept1* is overexpressed in ECs isolated from *Cept1^fl/fl^ Cre*^+^ mice (n=3). Gene expression of *Pparα, Acox1*, *Vegfa,* and *Vegf2r* is also provided (n=3). D) Western blotting demonstrates that CEPT1 content is increased in HUVECs transduced with *Cept1* lentiviral particles (MOI 5). Representative blots of VEGF2R, PPAR*α*, ACOX1, p-AKT, t-AKT and p-eNOS, t-eNOS are also provided. E) Quantitative representation of protein content relative to GAPDH as a loading control (n=3). F) RT-PCR analysis demonstrates *Cept1* is overexpressed in transduced ECs (n=3). Gene expression of *Pparα, Acox1*, *Vegfa,* and *Vegf2r* is also provided (n=3). Data are presented as mean ± SEM. * p<0.05; ** p<0.01; *** p<0.001. p values were from 1-way ANOVA and unpaired Student *t*-test.

### *Cept1* promotes the p-AKT/p-eNOS signaling in a *Pparα*-dependent manner

Since previous work has shown that CEPT1 can impact cell signaling in a PPARα-dependent fashion, we evaluated whether CEPT1-mediated EC activation is also PPARα-dependent. HUVECs were transduced with lentivirus to overexpress *Cept1* (p<0.05; Figure 6A) and were also transfected with si*Pparα* for efficient selective knockdown of *Pparα* (p<0.05; Figure 6B) in the same model. Selective knockdown of *Pparα* significantly reduced HUVEC cell migration (p<0.05; Figure 6C-D), and tubule formation (p<0.05; Figure 6E-F). Chemical inhibition with GW6471 (PPARα inhibitor), ZM323881 (VEGF2R inhibitor), LY294002 (AKT inhibitor), and L-*NAME* (eNOS inhibitor) also significantly reduced *Cept1*-induced migration in EC isolated from *Cept^fl/fl^ Cre*^+^ mice and *Cept1* overexpressed HUVEC (p<0.05; Figure 6G-J). These findings suggest that *Cept1* promotes EC angiogenic functions via VEGF-A, p-AKT, and p-eNOS signaling.

**Figure 6:**
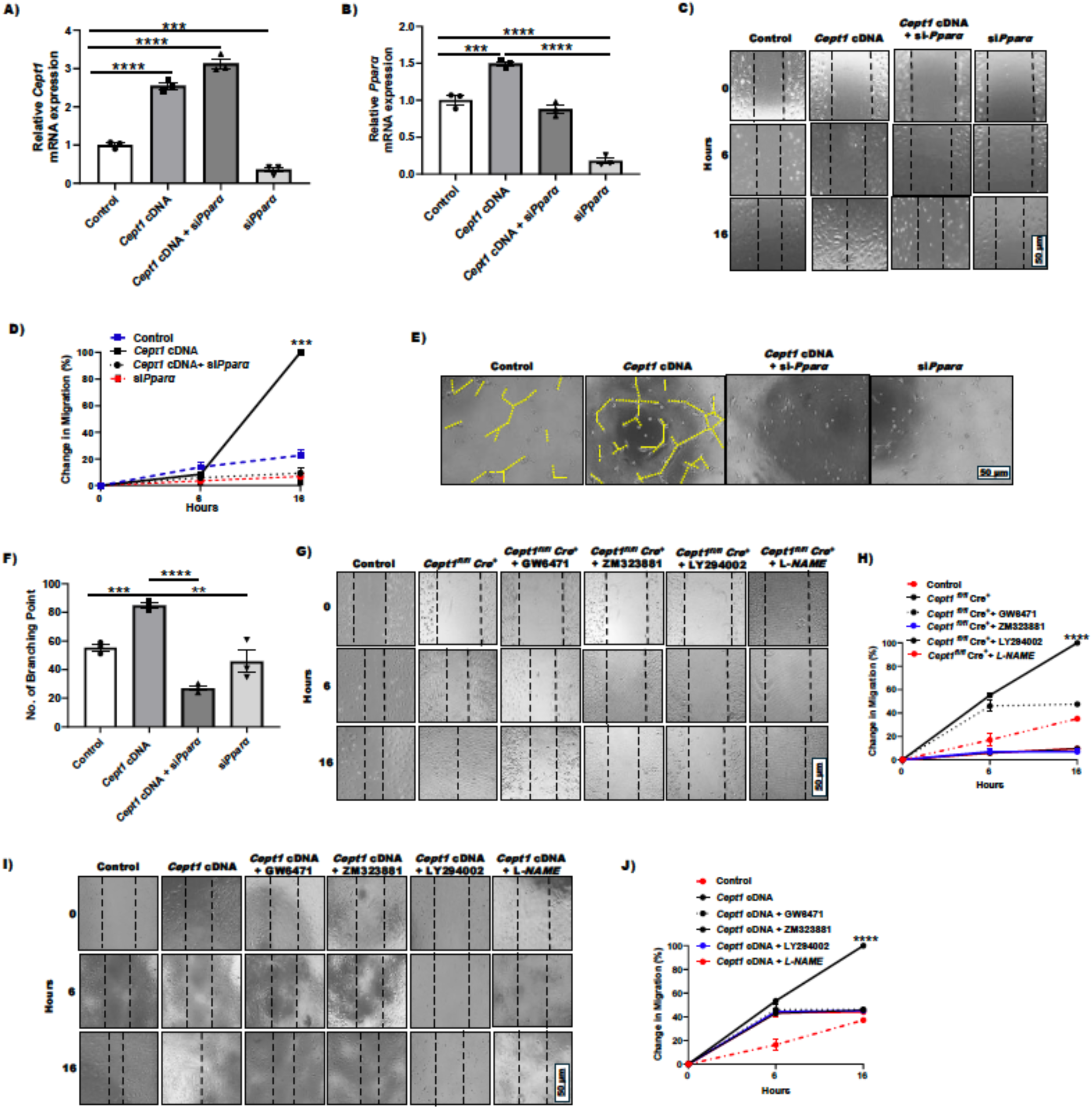
*Cept1* promotes *Vegfa, p-AKT,* and *p-eNOS* signaling pathways in a *Pparα*-dependent manner. A-B) HUVECs were transfected with siRNA targeting *Pparα* or siRNA control (si-Ct) for 72 h, and the expression of *Pparα* was confirmed through RT-PCR. Relative gene expression of *Cept1* and *Pparα* was performed using RT-PCR (n=3). C-D) Representative cell migration assay images at t= 0, 6, and 16h were taken from HUVEC *Cept1* cDNA, *Cept1* cDNA + si*Pparα*, and si*Pparα*. A quantitative graphical representation of cell migration was shown (n=3). E-F) Representative images and quantitative analysis of tubule formation of HUVEC with different conditions after 6h of incubation of Matrigel (n=3). G-H) EC from *Cept1^fl/fl^ Cre*^+^ were treated with GW6471 (10uM), ZM323881 (80nM), LY294002 (30uM), and L-*NAME* (0.5uM) and cell migration at t = 0, 6, and 16h was assessed and quantified, relative to EC from *Cept1^fl/fl^ Cre*^+^ and control (n=3). I-J) HUVEC were treated with GW6471 (10uM), ZM323881 (80nM), LY294002 (30uM), and L-*NAME* (0.5uM) and cell migration at t = 0, 6, and 16h was assessed and quantified, relative to control (n=3). Data are presented as mean ± SEM. * p<0.05; ** p<0.01. *P* values were from 1-way ANOVA and unpaired Student *t*-test.

## Discussion

Previously we observed that *Cept1* is essential for EC function and recovery following hind-limb ischemia.^17^ Here we expand on these findings and demonstrate that *Cept1* also induces molecular mechanisms responsible for enhanced recovery in the setting of ischemia. Specifically, we observed that CEPT1 has altered content in diseased peripheral arterial tissue in individuals with diabetes and PAD, and that this enzyme enhances EC function and viability. Additionally, we observed increased ACOX1 and VEGF2R content in the intimal of patients with PAD and diabetes. Similar trends were observed with conditional EC-specific *Cept1* overexpression in mice with STZ-induced diabetes. These mice had improved recovery, higher limb perfusion, and increased microvascular angiogenesis. *In vitro,* primary ECs and HUVECs that over-expressed *Cept1* also had enhanced EC function, and angiogenic signaling with increased p-Akt and p-eNOS. *Cept1*-induced EC function appeared to be highly dependent on p-Akt, p-eNOS, as well as PPARα. Collectively, these findings demonstrate that *Cept1* promotes angiogenesis and may play an important regenerative role in mitigating ischemic-induced tissue complications.

Knockdown of *Cept1* in ECs leads to decreased cellular function and ultimately reduces microvascular angiogenesis and tissue recovery following HLI.^17^ This phenotype is further exacerbated in the setting of STZ-induced diabetes. Here we explored whether endothelial *Cept1* overexpression can impact tissue recovery in the setting of ischemia and diabetes. Interestingly, we observed that STZ-treated *Cept1^fl/fl^ Cre*^+^ mice had increased hindlimb angiogenesis, and a more rapid improvement in perfusion and function. These findings highlight that increased CEPT1 can improve tissue perfusion and regeneration following ischemia in the setting of diabetes.

Our study used scRNA-seq to analyze >10,000 cells from murine aortic tissue. Notably, we identified five distinct clusters of cells representing the major primary cell types in the aorta: fibroblasts, VSMCs, EC1, EC2, and EC3. We observed three distinct EC subpopulations that highlighted their specific functional capabilities.^31^ *Cept1* was particularly overexpressed in the EC 2 cluster, which allowed us to conduct comparative analysis between the different EC clusters. ECs with increased *Cept1* expression also had increased expression of *Vegfa* and demonstrated enhanced molecular gene expression pathways that promote cell migration, angiogenesis, and wound healing.^34–36^ Conversely, inhibition of *Pparα* using GW6471 or selective knockdown reduced cell migration in ECs that overexpressed *Cept1.*^37,38^

It is widely acknowledged that diabetes represents a potent independent risk factor for PAD and increases the risk of complications such as lower-extremity foot wounds and amputations.^39–44^ We harvested peripheral arterial segments from patients with and without diabetesand observed that CEPT1 content was notably higher in Max-diseased arterial intima and that downstream PCs and PEs were also elevated. This is consistent with prior reports that lipid synthesis enzymes exhibit modified expression within the peripheral arterial intima of individuals with diabetes and advanced PAD.^17,45,46^

Our previous studies have investigated the role of *Cept1* and lipogenesis on EC function^17^ and implicated PPARα. This is consistent with prior reports that demonstrate that the CEPT1-derived PC 16:0/18:1 is a critical ligand that impacts *Pparα*-dependent gene expression, including expression of the β-fatty acid oxidation enzyme *Acox1*.^18^ A role for PPARα in angiogenesis has also been widely reported.^47–49^ WY14643 (PPARα-agonist) induces neovascularization in HUVECs and *in vivo* models^21^ consistent with a role for PPARα in angiogenesis. In our prior assessment of EC-specific knockout of *Cept1*, we observed a significant reduction in PPARα phosphorylation.^17^ We also observed that simultaneous knockdown of *Cept1* and *Pparα* severely impaired hind-limb perfusion and angiogenesis following unilateral femoral artery ligation. Interestingly, treatment of *Cept1* knockdown mice with fenofibrate (a PPARα agonist) improved hindlimb perfusion, but perfusion remained impaired in mice with both EC-specific *Cept1* and *Pparα* knockdown.^17^ These findings are consistent with our current observations that CEPT1-induced EC function and signaling is PPARα dependent.

To date, the largest prospective clinical trial to investigate the impact of a PPARα agonist in patients with type 2 diabetes is the Fenofibrate Intervention and Event Lowering in Diabetes (FIELD) study, which enrolled 9,795 patients from 63 clinical sites.^50^ Fenofibrate is a PPARα agonist that treats dyslipidemia.^51^ The study was consistent with some cardiovascular benefits, including reductions in microvascular complications, such as diabetic retinopathy and nephropathy.^52–58^ For lower extremity endpoints, the study found a reduction in lower-limb amputations, independent of lipid-lowering effects, suggesting a direct role in vascular health. While amputation was not a primary endpoint of the FIELD study, its findings suggest that PPARα agonists like Fenofibrate may have limb salvage benefits, potentially mitigating the morbidity and disability associated with type 2 diabetes and PAD.^58^ Our study supports this notion and provides molecular evidence that angiogenic EC function and signaling are dependent on PPARα activation, even in the setting of CEPT1 overexpression, further emphasizing the therapeutic potential of PPARα modulation in diabetic vascular disease.

It is known that *Pparα* stimulates angiogenesis by activating expression of *Vegfa* and *Vegfr2* genes.^21^ Zhang et. al. reported the induced release of pro-angiogenic factors in endothelial cells with the administration of WY-14 643 (a PPARα Agonist).^59^ In vivo, PPARα has been shown to promote angiogenesis in a VEGF-A-dependent manner, enhance nitric oxide (NO)-mediated vasodilation, and promote the release of p-AKT to protect against ischemia-induced tissue damage.^21,60^ PPARα can also promote differentiation of endothelial progenitor cells (EPCs) to become ECs, release VEGF-A, and activate downstream AKT signaling.^61^ Many have reported that this process is essential for tissue regeneration at sites of vascular injury and in the promotion of neovascularization.^62–64^ Inhibition of PPARα suppresses VEGF-A-induced angiogenesis *in vitro* and *in vivo*.^21^ *Ppara^-/-^*mice have blunted ischemia-induced angiogenesis and reduced VEGF content in ischemic tissue.^21^ In our study, we build upon these prior findings and demonstrate that CEPT1 appears to play an important upstream role in activating PPARα, and promoting VEGF-A signaling. Selective knockdown of *Pparα* using siRNA as well as pharmacological inhibition of PPARα with GW6471 corroborates prior study findings.^21^

In conclusion, we observed that CEPT1 overexpression in the endothelium can play an important role in activating EC function and angiogenic signaling. This signaling is dependent on PPARα and appears to play an important role in tissue regeneration and healing following ischemic insult - particularly in the setting of diabetes. Our findings confirm that PPARα activation can play an important role in the setting if peripheral limb ischemia and type 2 diabetes and suggest that further clinical investigation in this area is warranted to reduce the morbidity associated with PAD in this population.

## Notes

### Competing Interest Statement

The authors have declared no competing interest.

